# The age-regulated zinc finger factor ZNF367 is a new modulator of embryonic neurogenesis

**DOI:** 10.1101/255919

**Authors:** Valentina Naef, Sara Monticelli, Debora Corsinovi, Maria Teresa Mazzetto, Alessandro Cellerino, Michela Ori

## Abstract

Global population aging is one of the major social and economic challenges of contemporary society. During aging the progressive decline in physiological functions has serious consequences for all organs including brain. The age-related incidence of neurodegenerative diseases coincides with the sharp decline of the amount and functionality of adult neural stem cells. Recently, we identified a short list of brain age-regulated genes by means of next-generation sequencing. Among them *znf367* codes for a transcription factor that represents a central node in gene coregulation networks during aging but its function, in the central nervous system (CNS), is completely unknown. As proof of concept we analyzed the role of *znf367* during neurogenesis. By means of a gene loss of function approach limited to the CNS, we suggested that *znf367* might act as a key controller of the neuroblasts cell cycle, particularly in the progression of mitosis and spindle check-point. Using a candidate gene approach, based on a weighted-gene co-expression network analysis, we suggested possible targets of znf367 such as *fancd2* and *ska3*. The age-related decline of *znf36*7 well correlated with its role during embryonic neurogenesis opening new lines of investigation to improved maintenance and even repair of neuronal function.

## Introduction

The age-related incidence of many brain diseases coincides with a reduced adult neurogenic potential. The regenerative capability and the amount of adult neural stem cells (aNSCs) decline with age, contributing to the reduced functionality of the aged brain. Despite the great interest in age related diseases, the molecular factors responsible for age-dependent decay of aNSCs function and the transition between stemness and differentiating properties of these precursors are almost unknown. Recently, we identified a set of evolutionarily-conserved genes expressed in aNSCs and age-regulated by RNA-seq analysis in the short-lived fish *Nothobranchius furzeri*, a well-established animal model in aging studies. Among them, zinc finger protein 367 (znf367) was suggested to occupy a central position in a regulatory network controlling cell cycle progression and DNA replication. We found that *znf367* is expressed in the adult brain of *N. furzeri*, where its RNA level decreases with age, and in neuroblasts and retinoblasts of developing *Zebrafish* embryos ^1^. Znf367 belongs to the Zinc finger (ZNF) transcription factors family that represents a large class of proteins that are encoded by 2 % of human genes ^2, 3^. Their functions included DNA recognition, RNA packaging, transcriptional activation, regulation of apoptosis, protein folding and assembly, and lipid binding. Zinc finger proteins have an evolutionarily conserved structure and the ones containing the Cys_2_-His_2_ motif constitute the largest family. The function of the majority of ZNF genes is largely unknown, but some of them play a critical role in the development and differentiation of the nervous system. For instance, the Kruppel-like zinc finger transcription factor *Zic* has multiple roles in the regulation of proliferation and differentiation of neural progenitors in the medial forebrain and cerebellum ^4^. The *Ikaros* family of transcription factors is characterized by two sets of highly conserved Cys_2_His_2-_type zinc finger motif and is involved in the maturation and differentiation of striatal medium spiny neurons ^5^. *Znf367* gene (also known as *ZFF29*) has been initially isolated in human fetal liver erythroid cells. In the human genome, this gene is on chr 9q and two alterative mRNA splicing products, were identified and designated ZFF29a and ZFF29b. They both code for nuclear proteins, but only ZFF29b seems to act as an activator factor of erythroid gene promoters^6^. In Human SW13 adrenocortical carcinoma cell line, *znf367* is overexpressed and in this cell line *Znf367* downregulation caused an increase of cellular proliferation, invasion and migration^7^. Furthermore, *znf367* was also identified as a potential tissue-specific biomarker correlated with breast cancer where its expression level is dysregulated influencing cell proliferation, differentiation and metastatic processes^8^. To our knowledge, there are no data available regarding the putative role of *znf367* in the Central Nervous System (CNS) during embryonic and adult neurogenesis. In order to characterize the biological function of *znf367* in vertebrates CNS, we analyzed its role during *Xenopus laevis* neurogenesis. The clawed frog *Xenopus* is the gold standard as animal model to perform functional screening of genes. In *Xenopus*, it is possible to microinject mRNAs or morpholino oligos in just one side of the early cleaving embryo and compare, in every single embryo, the manipulated side of the embryo with its wild-type counterpart that represent a perfect internal control. This model gave us also the unique possibility among vertebrates, to rapidly perform gene loss of function experiments in a tissue specific manner thanks to the well-defined fate map of each blastomere of the early cleaving embryo. This allowed us to target specific *znf367* morpholinos to the central nervous system without interfering with the normal development of all other tissues. In this paper, we showed that *znf367* is expressed in the developing CNS in Xenopus and it could have a key role in the primary neurogenesis regulating the neuroblast progression of mitosis. This aspect well correlates with its gene expression decline during CNS aging, suggesting that *znf367* could represents a new key piece in the complex mosaic of developmental neurobiology and aging research.

## Results

### Evolutionary conservation and embryonic expression analysis of *znf367*

To verify the evolutionary conservation of znf367 sequence in vertebrates we performed an in silico analysis of the amino acid sequences of ZNF367 in *Homo sapiens* (both splicing variants: ZFF29a and ZFF29b), *Nothobranchius furzeri* and *Xenopus laevis* (both splicing variants: *znf367a* and *znf367b*). This approach revealed a high conservation of znf367 with a 66% of identity between the human and Xenopus aminoacidic sequence that reached the 98% at the level of the zinc finger domains (Fig 1) suggesting a conserved putative znf367 function in vertebrates, from fish to tetrapods and primates. To analyze the spatio-temporal gene expression pattern of *znf367*, whole mount in situ hybridization (WISH) was performed on xenopus embryos at different stages. Z*nf367* is expressed maternally in the animal pole in Xenopus embryos at blastula stage (Fig. 2A-B) when compared to sense control probe treated siblings (Fig. 2A). At neurula stage *znf367* is expressed in the neural tube, in the eye fields, in the pre-placodal territory and in the neural crest cells (NCC) (Fig. 2C). At tadpole stage *znf367* is widely expressed in the central nervous system, in the eye, in otic vesicle and in the NCC migrated in the branchial pouches (Fig. 2D). At larval stages of development, *znf367* is still widely expressed in the CNS as showed in transverse sections (Fig. 2E-F)

**Figure 1.**
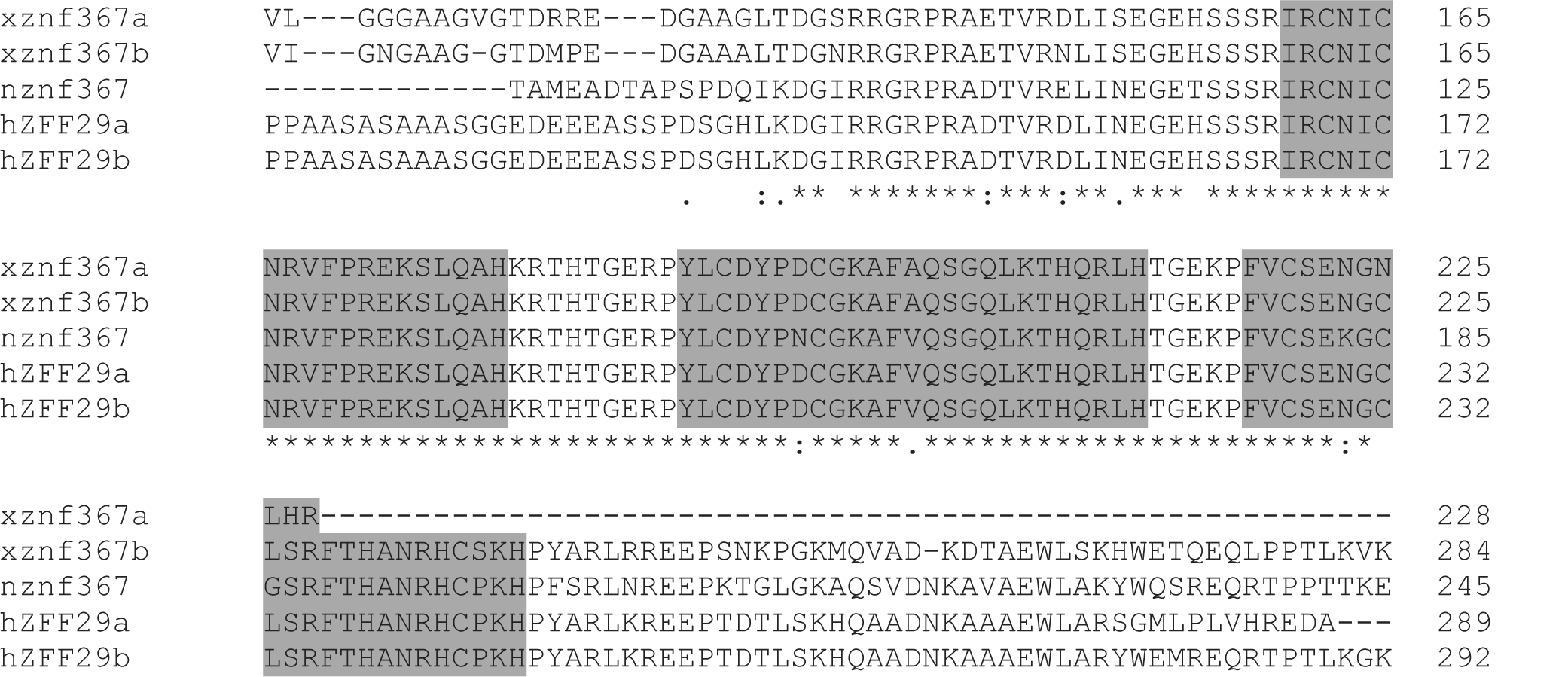
Multiple sequence alignments of *znf367* amino acid sequences containing the zinc finger domains of *znf367* from *X. laevis* (both splicing variants *znf367a* and *znf367b*) and human (both splicing variants ZFF29a and ZFF29b) and *N.furzeri* (n) were performed using Clustal Omega. The conserved zinc-binding domains are highlighted in gray.

**Figure 2.**
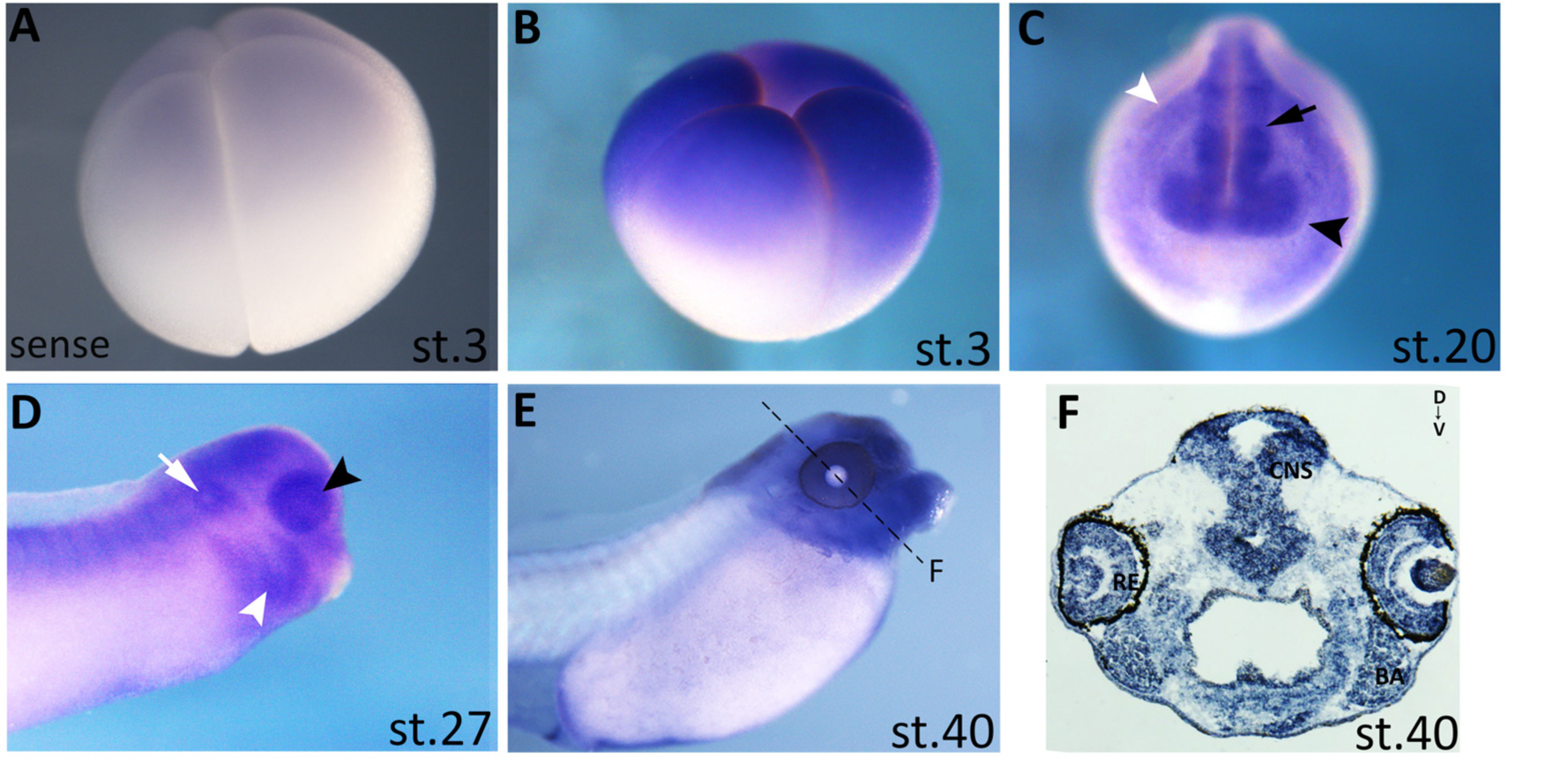
*Znf367* gene expression pattern during *Xenopus laevis* development. **(A-B)** *Znf367* expression at blastula stage (stage 3) using sense control probe (A) and antisense probe (B). **(C-D)** At neurula (C) and at tadpole stages (D) *znf367* is expressed in the neural tube (black arrow), the developing eye (black arrowed), the neural crest cells (white arrowed) in the otic vesicle (white arrow) and in the most anterior regions of the nervous system **(E-F)** Stage 40 embryo is shown in lateral view (E) and in a transversal section at the level of the hindbrain (F). CNS, central nervous system; BA, branchial arches.

### *Znf367* Knockdown inhibits neuronal differentiation in *Xenopus laevis* embryos

To investigate the *znf367* function during neurogenesis in *Xenopus*, we performed knockdown experiment using a specific antisense oligonucleotide morpholino designed to block the translation of the endogenous mRNA (ZNF367-MO). For all the experiments here described, injections were performed unilaterally into one dorso-animal blastomere at four cells stage to target the neural tissue. The un-injected side served as internal control and the co-injection of 250 pg *gfp* RNA was used to screen the embryos correctly injected (Fig. 3A). The standard Gene Tools Control-morpholino (co-MO) was used to control for non-specific embryo responses. At neurula stage (st.18), when the neural tube is just closed, *znf367* morphants showed a strong reduction of post-mitotic neurons expressing *N-tubulin* and *elrC* (also known as *HuC)* on the injected side of the embryos compared to the control side and the co-MO injected embryos (Fig. 3B-3E’). These data are confirmed also by qRT-PCR analysis that showed a significant reduction of both neuronal markers in *znf367* morphants (Fig. 3F-G). Interestingly, the injection of ZNF367-MO did not affect the expression of *ngnr1,* a proneural marker necessary for the specification of primary neurons ^19^(n=53) (Fig. 3H-H’) suggesting a role of *znf367* during neuronal differentiation but not in neuronal specification. The lack of differentiated primary neurons in *znf367* morphants could be due to an increase in cell apoptosis during the differentiation process. To evaluate this aspect, we performed a TUNEL assay in *znf367* morphants at neurula stage. TUNEL staining revealed not-significant increase in TUNEL positive cells between the *znf367* injected side and the un-injected control side of each analyzed embryo (Fig. 3I).

**Figure 3.**
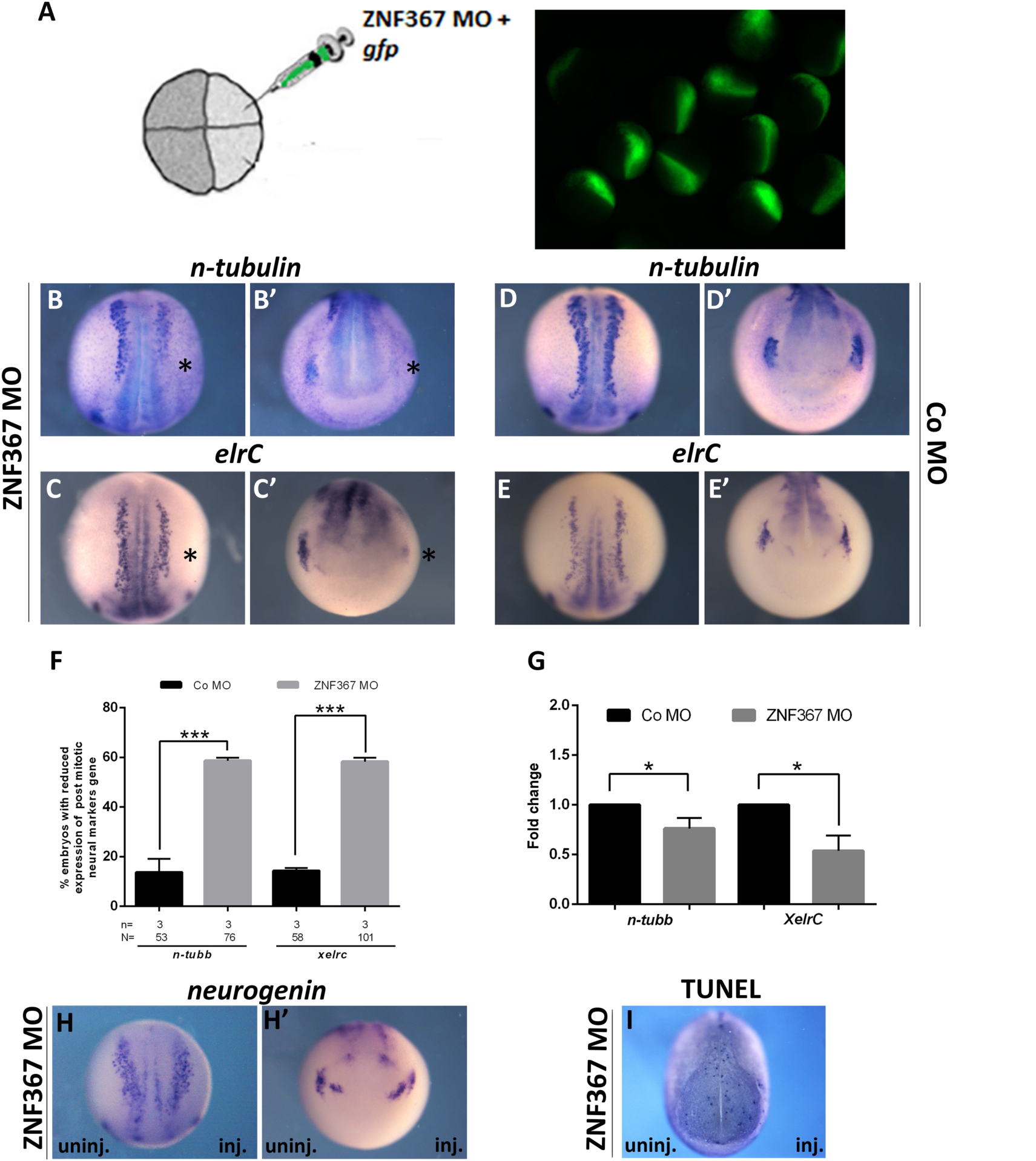
**(A)** Embryos injected with *gfp (*250 pg) and either ZNF367-MO or Co-MO (9 ng) at one dorsal blastomere at the four-cells stage showed fluorescence only in the neural plate at neurula stage (st. 18). In each panel, the injected side of the embryo is indicated by the asterisk. **(B-E’)** mRNA distribution of *N-tubulin* and *elrC* in *znf367* morphants and controls. (B, C) dorsal view and (B’, C’) frontal view of neurula morphants showed a clear down regulation of primary neurons markers. (D, E) dorsal view and (D’, E’) frontal view of neurula control embryos. **(F)** Quantification of the data in A and B **(G)** qRT-PCR analysis. Relative expression levels of each gene are normalized to Glyceraldehyde 3-phosphate dehydrogenase (*gapdh*) **(H-H’** *; embryos are shown in a dorsal view anterior down and in an anterior view*) mRNA distribution of *neurogenin* in *znf367* morphants. **(I;** *embryo is shown in an anterior-dorsal view*) TUNEL staining. ZNF367-MO injection does not lead to an increase of TUNEL positive cells compare to the un-injected side. Abbreviations: n, number of independent experiments; N, number of evaluate embryos in total; Error bars indicate standard error of the means (s.e.m); * p≤0,05*** p≤0,001. P-value were calculated by Student’s t-test.

### *Znf367* knockdown increases proliferation markers in *Xenopus laevis* embryos

To determine, whether the observed loss of post-mitotic neurons in *znf367* morphants was the consequence of impairment in the maintenance of the neuronal progenitors pool, we examined the expression of the stemness genes *sox2* and *rx1*, in the injected embryos. *Sox2* and *rx1* are involved in maintaining neuroblasts and retinoblasts as cycling precursors in the neural plate ^20,21^. The expression domains of *sox2* and *rx1* were expanded on the ZNF367-MO injected side of the embryo as compared to either the un-injected and Co-Mo injected sides (Fig. 4A-B). These data were confirmed also by qRT-PCR analysis that showed a significant increase of both mRNAs in *znf367* morphants (Fig. 3B). On the base of these results we can suggest that the *znf367* knockdown enhances self-renewal at the expenses of differentiation. For testing the specificity of the ZNF367-MO to induce this phenotype, we performed functional rescue experiments by co-injecting 9 ng ZNF367 MO together with 500 pg full-length *Xenopus znf367* mRNA. We observed a restoration of the phenotype of the injected embryos visualized by the *sox2* and *elrc* markers at neurula stage (Fig. 20) (30% of rescue for *sox2* n=114; 25% of rescue for *elrc* n= 100) (Fig. 4D). To further verify whether *znf367* downregulation could alter the regulation of neuroblasts proliferation, we also examined the mRNA expression of *pcna (proliferating cell nuclear antigen)* and we directly counted mitotically active cells marked by antibody specific for phosphorylated H3 (p-H3). *Znf367* morphants showed an increased *pcna* mRNA expression both in WISH (Fig. 4E-F) and qRT-PCR experiments (Fig. 4G). p-H3 staining showed a significant increase in mitotic cells number upon ZNF367-MO injection as compared to the control side (Fig. 4H-I). Given that a larger pool of neuroblasts did not correspond to an increased number of differentiated cells in the absence of apoptotic cell death, it is tempting to speculate that *znf367* could be required to exit the M phase or in the control of the mitotic check point that precedes the anaphase. To test this hypothesis, we first evaluated the relative expression of *cyclin B1* that is expressed predominantly during M phase of the cell cycle^22^, by qRT-PCR analysis of *znf367* morphants. This experiment revealed a significant increase of *cyclin B1* expression in znf367 morphants (Fig 4L) indicating again that *znf367* deficient neuroblasts could enter in M phase but they could not correctly exit mitosis and differentiate. The differentiation of neuronal progenitors requires the withdrawal from the cell cycle, driven by cell cycle inhibitors such as *pak3 (p21)* and *p27* ^23^; ^24^. Coherently with the increase in mitotically active cells, a mild loss of *p27* expression (phenotype 55%, n=93) was observed in neurula morphants indicating that *znf367* depleted neuroblasts are unable to exit cell cycle (Fig. 4M-N). These data let us to hypothesize that *znf367* could be involved in the cell cycle exit and/or for the initiation of maintenance of a differentiated state. Finally, we examined morphants at tailbud stage by performing WHIS using *rx1* and *elrD* (also known as *HuD*). *ElrD* labeled post mitotic neurons in the neural tube and the developing cranial ganglia^25^. As stated above, *znf367* knockdown, but not control MO, caused an increase in *rx1* gene expression (phenotype 52%, n=72) (Fig. 4O) while inhibited neuronal differentiation affecting the expression of *elrD* (phenotype 54% n=64) (Fig. 4P). These data showed that the effects of ZNF367 depletion are not recovered even in the late phases of primary neurogenesis.

**Figure 4.**
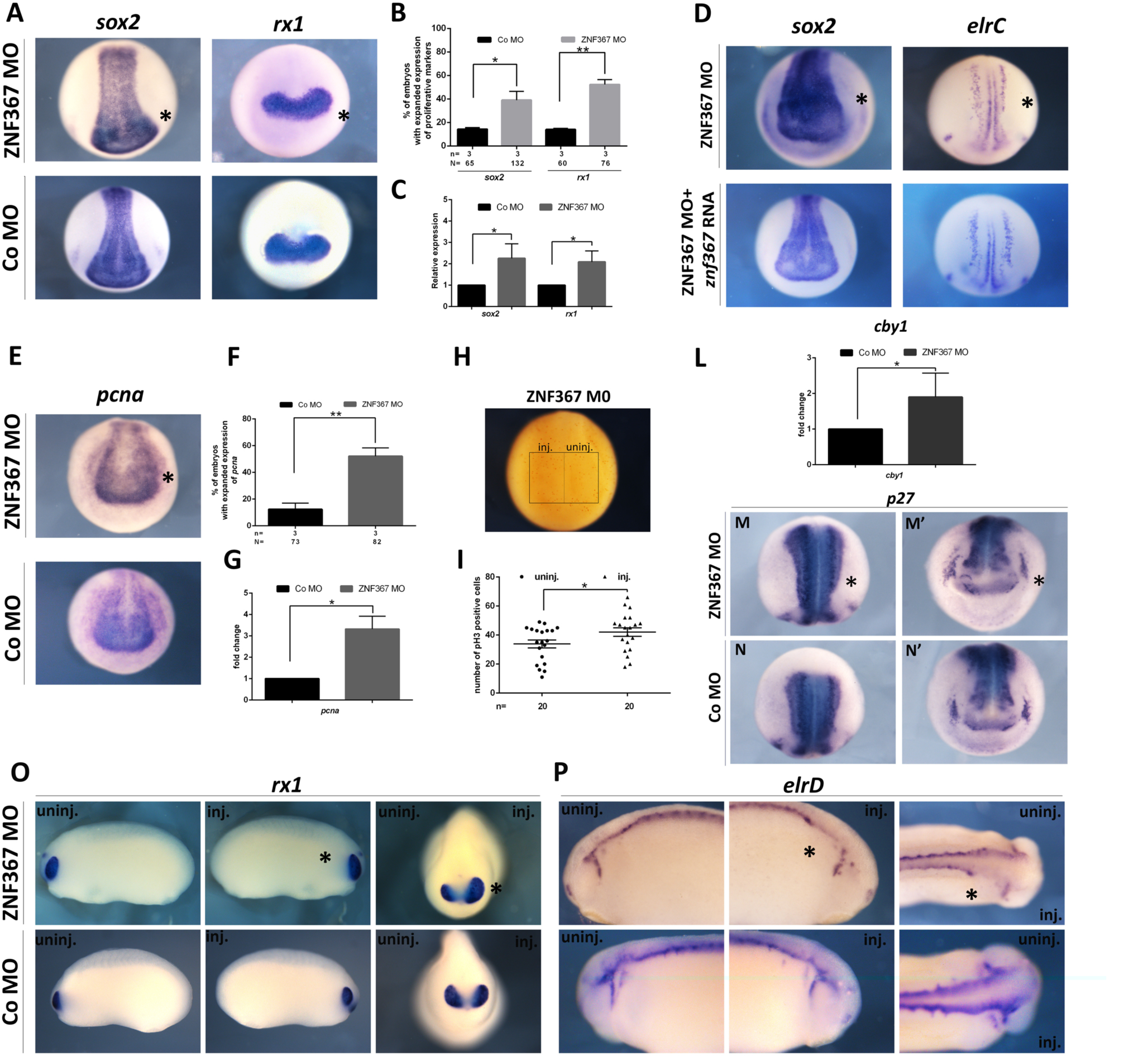
In each panel, the asterisk indicates the injected side of the embryo. **(A)** mRNA distribution of *sox2* and *rx1* in *znf367* morphants and controls. **(B)** Statistical analysis of the data in A. (**C**) qRT-PCR analysis. Relative expression levels of each gene are normalized to *glyceraldehyde 3-phosphate dehydrogenase* (*gapdh*). **(D)** The morphants phenotype can be rescued by the co-injecting morpholino plus full length Xenopus *znf367* mRNA as shown by the recovered expression of *sox2* and *elrC*. **(E)** mRNA distribution of *pcna* in *znf367* morphants and control. **(F)** Statistical analysis of the data in E. **(G)** RT-PCR analysis revealed a significant increase of *pcna*. Relative expression levels of gene are normalized to *glyceraldehyde 3-phosphate dehydrogenase* (*gapdh*). **(H-I)** *Znf367* depletion leads to a significant reduction of proliferating cells compared to the un-injected side. pH3 positive cells were counted in the areas defined by the black rectangles. Statistical evaluation of the data is shown (I), and p-value was calculated by a nonparametric, one-tailed Mann-Whitney rank sum test. **(L)** RT-PCR analysis of *cyclinB1* (*cby1*). Relative expression level of gene is normalized to *glyceraldehyde 3-phosphate dehydrogenase* (*gadph*). **(M-M’)** *p27* is down regulated at stage 18 in *znf367* morphants. (M) dorsal view; (M’) frontal view. **(N-N’).** *p27* expression in control embryo, (N) dorsal view; (N’) frontal view. **(0-P)** *rx1* and *elrD* expression at tailbud stages confirmed the phenotype observed at neurula stages: increased expression of a proliferation marker (*rx1*) and downregulation of neuronal differentiation marker (*elrD*). Abbreviations: n, number of independent experiments; N, number of evaluated embryos in total; Error bars indicate standard error of the means (s.e.m); * p≤0,05;** p≤0,01. P-value were calculated by Student’s t-test.

### Identification of putative Z*nf367* targets: a candidate gene approach

Our previous results suggested that *znf367* could represent a hub in the control of gene expression in the *N. furzeri* brain. In order to test the conservation of this co-regulation across species, we analyzed CORTECON^18^ a public dataset of RNA-seq during cortical differentiation of human embryonic stem cells (hESCs) using weighted-gene co-expression network analysis (WGCNA)^17^. WGCNA constructs co-expression networks based on topological criteria, it was shown to be more robust than simple correlation and it has become the method of choice for gene expression studies in the nervous system. We therefore tested the conservation of gene co-expression networks between *N. furzeri* brain and human neuronal differentiation *in vitro.* We identified a conserved module that contains *znf367* (Fig. 5). Then, we tested whether *znf367* can be considered a hub in both species by computing its connectivity. *Znf367* was among the top connected genes in the gene module in both species (98% percentile in *N. furzeri* and 92% percentile in human cells). Gene Ontology overrepresentation analysis revealed that cell-cycle related terms are highly overrepresented in this module. It should be also noted that all these genes have high expression in the hESCs, they are down-regulated during early differentiation and show a peak of expression around 12 days of differentiation *in vitro* that correspond to the period of cortical specification^18^. Among the genes that showed the highest topological overlap, we particularly noted an enrichment in genes known to be involved in the progression of mitosis and in the mitotic spindle check point (Fig.5B). This corroborate the idea that *znf367* had a role in the control of cell cycle and it could be preeminent in mitosis, when the diving cell has the fundamental task to correctly arrange the genetic content in the two daughter cells. To verify our hypothesis, we decided to test by qRT-PCR the expression of three of the closest genes to *znf367* showed in the network (Fig. 6). We analyzed the expression level of *smc2*, *ska3* and *fancd2* in *znf367* morphants. *Smc2* gene codes for one of the condensin part of the Structural Maintenance of Chromosomes (SMC) protein complexes, which play key roles in the regulation of higher-order chromosome organization and its role is crucial in the chromatin compaction in the prophase ^26^. *Ska3* is one of the spindle and kinetochore-associated (Ska) proteins required for accurate chromosome segregation during mitosis. During mitosis the cyclin-dependent kinase Cdk1 phosphorylates *Ska3* to promote its direct binding to the Ndc80 complex, (also present in the Znf367 network). This event is required for the overcoming of spindle checkpoint and the beginning of anaphase ^27^, ^28^. *Fancd2* encodes for a nuclear effector protein that is monoubiquitinated in response to DNA damage, targeting it to nuclear foci where it preserves chromosomal integrity. Mutations in the Fanc family are causative of the Fanconi Anemia in humans. Greater than 60% of Fanconi anemia patients have developmental defects, such as growth retardation, short stature, microcephaly, and microphthalmia at birth, in addition to a highly elevated risk of bone marrow failure in the first decade of life. This gene draws our attention as its knock down in zebrafish embryos induced microcephaly, microphthalmia and pericardial edema ^29^. It has been demonstrated that this factor is crucial for the S-phase rescue of damaged DNA but also for the safeguarding of chromosome stability during mitosis^30,31^. The results obtained in three independent experiments, showed a significant increase in *fancd2* and *ska3* gene expression in *znf367* morphants. The *smc2* gene expression level followed the same trend without a statistical significance. These results seem to suggest that *znf367* could have a major role in the control of chromosome stability and the functionality of the spindle check point. The loss of *znf367* gene function could alter the strict control on the cell cycle progression during metaphase causing the expansion of the neuroblasts and retinoblasts territories and the increase of mitotic cell numbers observed in *znf367* morphants. It is also interesting to note that in human tumor cells the overexpression of FANCD2 and SKA3 are correlated with cancer cell proliferation maintenance and malignant transformation^32 33^. On the basis of our results it is also possible to hypothesize that *znf367* could act, in the control of the cell cycle, as a transcriptional repressor to allow the progression between metaphase and anaphase and the subsequent cell cycle exit and neuronal differentiation.

**Figure 5.**
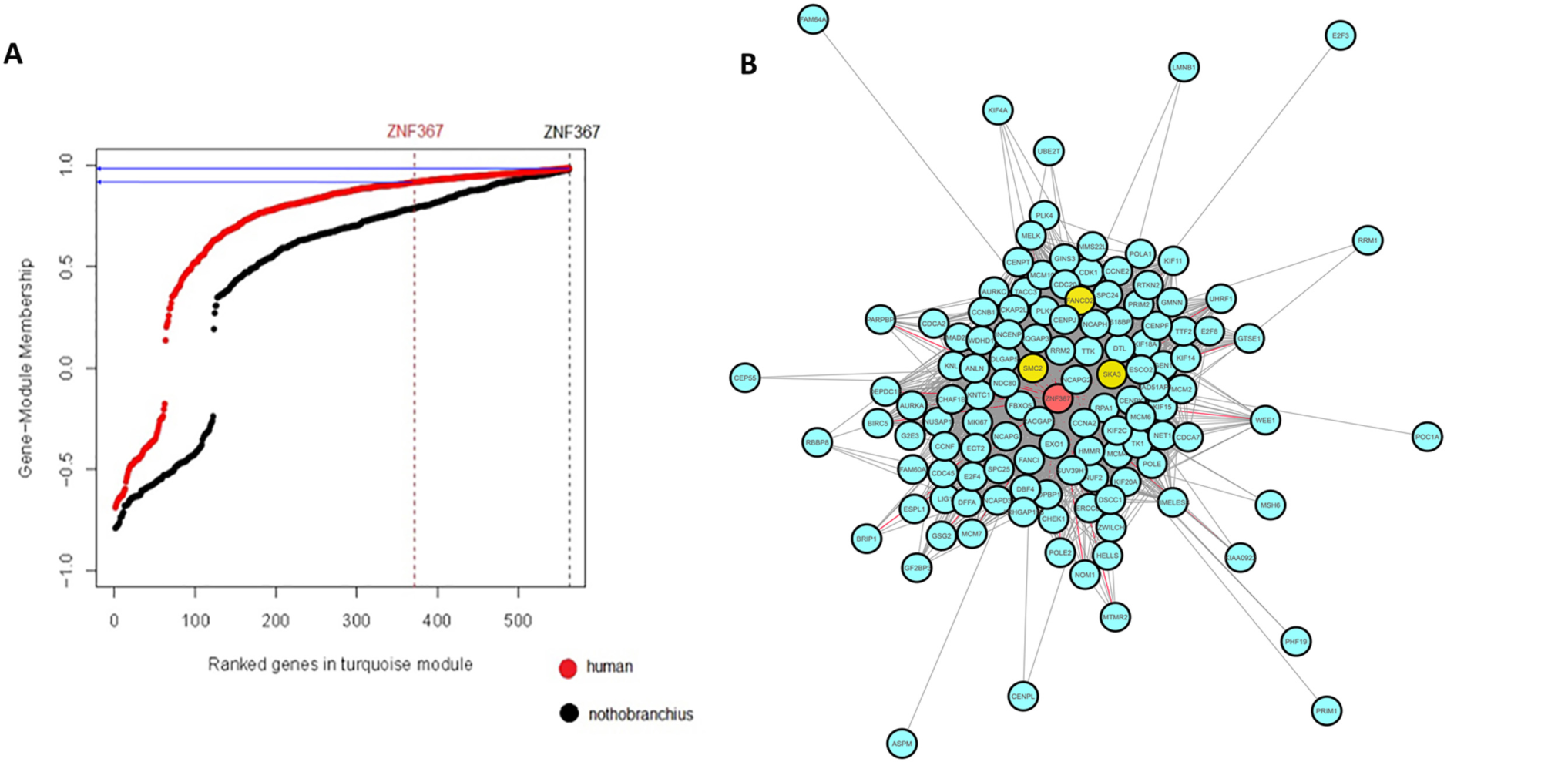
**(A)** Distribution of membership values for the “turquoise” module in human cells and N. furzeri brain. The vertical line indicates the rank of ZNF367 and the horizontal line its membership value (black for *N. furzeri* and red for *H. sapiens*). Please note due to the high convexity of the human distribution ZNF367 has very high membership despite its rank. **(B)** Central part of the gene module containing ZNF367. Only genes showing topological overlap > 0.3 are shown. ZNF367 is in red, while other genes used for further analysis (among ZNF367 neighbors) are in yellow (SMC2, SKA3 and FANCD2).

**Figure 6.**
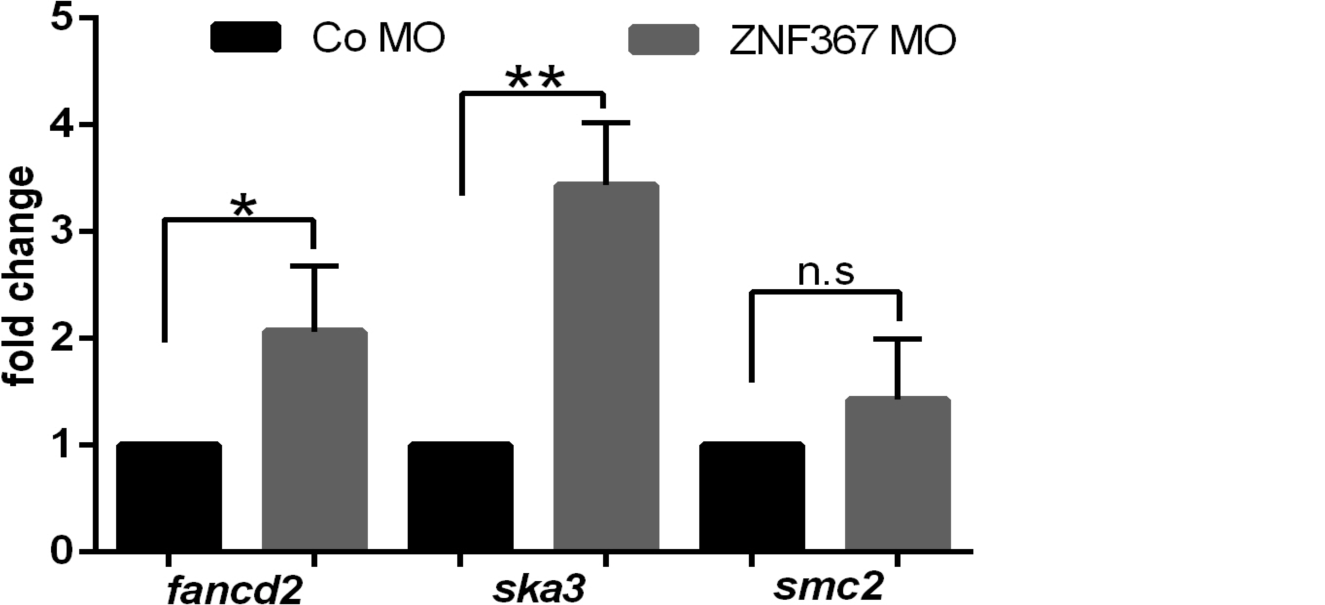
qRT-PCR analysis of *fancd2*, *ska3* and *smc2.* For each gene the relative expression level is normalized to *glyceraldehyde 3-phosphate dehydrogenase* (*gapdh*). Abbreviations: Error bars indicate standard error of the means (s.e.m); n.s., not significant. * p≤0,05;** p≤0,01. P-value were calculated by Student’s t-test

## Discussion

Aging is an inevitable and extremely complex, multifactorial process. It is characterised by the progressive physiological decline of organs and tissues linked to a reduction of their regenerative capabilities and the progressive exhaustion of adult neural stem cells. Interventions designed to target the underlying mechanisms of aging are expected to provide great benefit to human health and quality of life for elderly people. To provide a contribution to the field, we identified a short list of brain age-regulated genes of possible regulatory function, specifically associated with aNSCs in *N.furzeri,* an innovative animal model system in aging studies, by means of next-generation sequencing (NGS). These potential neurogenic regulators are down-regulated with age in an evolutionarily conserved manner and are also expressed in at least one neurogenic region of the zebrafish embryo. Among them, we found the zinc finger protein znf367 to be particularly intriguing. Network analysis identified *znf367* as a central node in gene co-regulation networks controlling cell cycle progression and DNA replication^1^. However, nothing is known about its role in the vertebrate nervous system. Our first aim was to provide an in vivo validation of the potential role of *znf367* during neurogenesis. We choose *Xenopus laevis* as a model system for the possibility to directly modulate the *znf367* expression in the CNS without affecting other tissues and to unveil its role in tetrapods. *Znf367* is expressed in the neural tissue of the early *Xenopus laevis* embryo including the eye field and in the neural crest cells. The spatial expression pattern suggested a role in the context of primary neurogenesis. This was further supported by the marked loss of post mitotic neurons upon knockdown of *znf367,* suggesting that *znf367* could be essential for neuronal differentiation. In Xenopus znf367 morphants, we did not observe an increase in apoptosis rate suggesting that the loss of post-mitotic neurons was not due to unspecific morpholino toxicity or to a specific triggering of apoptotic pathways. Indeed, we found that the loss of *znf367* function led to an increased expression of genes involved in the maintenance of neuroblasts and retinoblasts as proliferating precursors. Accordingly, we observed a significant increase in the number mitotic cells in *znf367* morphants. The intricate balance between proliferation and differentiation is of fundamental importance in development of the central nervous system. At early developmental stages, a period of extensive proliferation is needed to generate the required number of progenitor cells for correct tissue and organ formation, accompanied or closely followed by differentiation. After the closure of the neural tube, the epithelial lining of the ventricles become specialized, consisting of a single sheet of progenitor cells (neuroepithelial cells). These cells undergo symmetrical cell divisions during the proliferative period to self-renew and expand the pool of progenitors ^34^. The subsequent transition from a proliferative neural precursor cell to a post-mitotic neurons is a highly regulated step, which, in many instances, has been shown to involve a cascade of transcription factors that is triggered by pro-neural genes^35^. Differentiation of neural progenitor cells requires withdrawal from the cell cycle, which is regulated by the expression of cell cycle inhibitors such as *p27* in Xenopus^24^. Consistent with the increase in mitotically active cells, a reduced *p27* expression was observed in *znf367* morphants, thus raising the possibility that the neural progenitors are prevented from undergoing differentiation because they are not able to exit the cell cycle, remaining in an undifferentiated state. *Znf367* morphants also expressed high levels of *cyclin B1,* which is required to drive cells into mitotic division but that must be degraded to allow the anaphase. Again, this datum corroborates our hypothesis that *znf367* deficient neuroblasts could enter in M phase, but they could not correctly exit mitosis and differentiate. Given the requirement of *znf367* for both proliferation and neuronal differentiation of neuroectodermal cells, it is plausible that *znf367* could be required to exit the M phase or in the control of the spindle check point that precedes the anaphase. To have a wider view on the molecular mechanisms potentially regulated by *znf367,* we performed a correlation-based network analysis testing the conservation of gene co-expression networks between *N. furzeri* brain and human neuronal differentiation in vitro, identifying a conserved module that contains *znf367*. We noted enrichment in genes involved in the regulation of the cell cycle and specifically in the progression of mitosis and mitotic spindle check point. The involvement of *znf367* in the control of cell cycle is supported by functional studies that demonstrated its implication in regulating different aspects of cancer progression^7^. Some of the genes, correlated to *znf367* in our correlation analysis are, in fact, both implicated in CNS development as well as in cancer initiation and/or progression. Among these, we decided to closely analyze the relation between *znf367* and *smc2*, *ska3* and *fancd2*. Smc2 is part of the condensing complex required for the structural and functional organization of chromosomes^36^. Its role is crucial in the chromatin compaction in the prophase ^26^. In our functional study the loss of *znf367* seemed to interfere with *smc2* mRNA level but even if *smc2* mRNA seemed to be more abundant in *znf367* morphants in respect to controls, the results are suggestive of a trend but not statistically significant. *Fancd2* is essential during zebrafish CNS development to prevent neural cell apoptosis during neuroblasts proliferative expansion^29^. *Fancd2*, when mutated, is one of the causative gene of Fanconi anemia, an inherited disorder characterized by developmental defects, progressive bone marrow failure, and predisposition to cancer. In particular, *fancd2*, in postnatal and adult life is required for the functional maintenance of the hematopoietic stem cells pool^37^. The link between *znf367* and *fancd2* seems therefore particularly intriguing since the *znf367* function seemed to be required to repress *fancd2* expression and allow cells to inactivate the spindle checkpoint and proceed towards differentiation. The level of *fancd2* mRNA is, in fact, significantly up regulated in *znf367* morphants. It is tempting to speculate that during primary neurogenesis in Xenopus *znf367* could regulate *fancd2* expression level in order to define the pool of neuroblasts and coordinate the cell cycle exit necessary for the post-mitotic differentiation. *Ska3* is one of the spindle assembly checkpoint proteins. SKA3 is strongly associated to the kinetochores during prometaphase and metaphase while diminished its concentration during anaphase and it was lost telophase ^27^. Its major role is to contribute to the silencing of spindle checkpoint during metaphase and to the maintenance of chromosome cohesion in mitosis ^27^. In our *znf367* morphants, *ska3* is upregulated supporting the idea that *znf367* could play a key role in the control of mitosis and in particular during metaphase. As *ska3* has to be downregulate to allow the progression towards anaphase and telophase, the loss function of *znf367* could maintain abnormal high level of *ska3* keeping cells blocked in mitosis (metaphase). This analysis provided us a deeper view of the possible action of *znf367* during neurogenesis. Functional analysis on *znf367* morphants clearly pointed to a role of *znf367* in the control of the cell cycle and in the formation/maintenance of the neuroblasts pool. The loss of *znf367* caused a differentiation failure keeping an enlarged number of neuroblasts in mitosis. In this condition, neuroblasts did not activate an apoptotic pathway that can be prevented when *fancd2* is present^29^. The gene network analysis also suggested a possible function of *znf367* in the regulation of the spindle checkpoint during the metaphase acting on the expression level of *ska3*.

In conclusion, we unveiled a role for *znf367* during neurogenesis in vertebrates. In particular, *znf367* emerged as a key controller of the neuroblasts cell cycle and it seemed to act regulating the events that are strictly controlled during the metaphase to allow the progression of the cell cycle and the onset of anaphase. The observed age-related regulation of *znf36*7 well correlated with its role during embryonic neurogenesis giving a proof of concept of the continuity of molecular control in developing and adult neurogenesis. It will be of interest for future studies to identify both the upstream regulators and the downstream effectors of *znf367.* This is important not only due to the requirement of *znf367* during *X. laevis* neurogenesis but more generally for the identification of the molecular factors that allow better monitoring of stem cell renewal and differentiation. Our findings could represent the first step in defining new strategies to increase adult neurogenesis, leading to improved maintenance and even repair of neuronal function.

## Methods

### Synteny analysis of znf367

Synteny analysis was performed using the NCBI GeneBank for the following organisms: *Xenopus laevis znf367*a (NP_001085362.1); *Xenopus laevis znf367b* (XP_018114684 PREDICTED); *Homo sapiens ZFF29A* (AY554164.1) and *Homo sapiens ZFF29b* (AY554165.1); *Nothobranchius furzeri* (HADY01011608.1).

### Embryo preparation

Animal handling and care were performed in strict compliance with protocols approved by Italian Ministry of Public Health and of the local Ethical Committee of University of Pisa (authorization n.99/2012-A, 19.04.2012). *Xenopus laevis* embryos were obtained by hormone-induced laying and in vitro fertilization then reared in 0.1 X Marc’s Modified Ringer’s Solution (MMR 1×: 0.1 M NaCl, 2 mM KCl, 1 mM MgCl2, 5 mM HEPES pH 7.5) till the desired stage according to Nieuwkoop and Faber^9^.

### Morpholino oligonucleotides, cloning and microinjections

ZNF367 antisense Morpholino oligonucleotides (MO) and a standard Control MO were provided by Gene Tools, Philomath, OR, USA. ZNF367 MO sequence:5’-CAGCCTATCTGACATTTGTTACTAC-3’. Co MO sequence: 5’-CCTCTTACCTCAGTTACAATTTATA-3’. Microinjections were performed as described previously ^10^. Injected MO amounts were: 9 ng ZNF367 MO and 9 ng Control MO. Correct injections were verified by co-injected of 250 pg of GFP mRNA and using a fluorescence microscope. The un-injected side represents an internal control in each embryo. For functional rescue experiments, the open reading frame of *X. laevis znf367* (XM_018259195.1 PREDICTED) was cloned into the pCS2+. For Rescue experiments, 9 ng ZNF367 MO and 500 pg full-length *znf367* mRNA were co-injected. Capped *znf367* mRNA was obtained using the MegaScript in vitro transcription kit (Ambion), according to manufacturer’s instructions.

### In situ hybridization (ISH) experiments

Whole mount in situ hybridization (WISH) approaches was performed as described ^11^. BM purple (Roche) was used as a substrate for the alkaline phosphatase; digoxigenin-11-UTP-labelled sense and antisense RNA probes were generated via in vitro transcription. After color development embryos were post-fixed and bleached over light to remove the pigment. For ISH on cryosections (12 µm), embryos were fixed in 4% paraformaldehyde (PFA) in PBS, cryoprotected with 30% sucrose in PBS and embedded in Tissue-Tek O.C.T. compound (Sakura, 4583). ISH on cryosections was performed as described ^11^. The following plasmids were used for preparation of antisense RNA probes, enzyme used for linearization and polymerases are indicated: *X. laevis Znf367* EST clone image (ID_6637026) was cloned in pBKS-(XhoI, T7); *Pcna-pBSK* (SalI,T7); *sox2-pCS2+* (EcoR1,T7); *N-tubulin-pBKS*(NotI,T3); *elrC-pBKS* (NOTI,T7); *elrD-pBSK* (XhoI,T3); *rx1* ^12^. *nrg1-pBKS* (BamHI, T3); *p27-Pbsk* (BamHI, T7). Digoxigenin labelled sense RNA probe was generated for *znf367-pBKS-* (SacI; T3).

### TUNEL and PH3 staining in *Xenopus*

TUNEL (TdT-mediated dUTP-dig nick end labeling) and PH3 (phosho histone 3) staining was performed at neurula stage according to established protocols ^13 10^. TUNEL and PH3 positive cells were counted within defined areas in control and injected side of each manipulated embryo.

### Quantitative Reverse Transcription Polymerase Chain Reaction (qRT-PCR)

Total RNA was extracted from 15 Xenopus morphants using Nucleospin^®^ RNA (Macherey-Nagel) according to the manufacturer’s instruction. cDNA was prepared by using iScript™ cDNA Synthesis Kit (Bio-Rad) and quantitative real-time PCR was performed using GoTaq^®^qPCR master mix (Promega) according to the manufacturer’s instruction. Relative expression levels of each gene were calculated using the 2^−ΔΔCt^ method ^14^ and normalized to glyceraldehyde 3-phosphate dehydrogenase (GAPDH). The following primers were used to perform qRT-PCR: *pcna* ^15^; *N-tubulin* and *sox2* (De Robertis’s lab, web site: http://www.hhmi.ucla.edu/derobertis/); *elrC* ^16^; *cby1* (Forward: 5’-TGAAGCGGTTCCAGTTGTCG-3’; Reverse: 5’-TTGGTGGCAACAACCCTCTT-3’); *ska3* (Forward: 5’-ACCGGAACTTTCCTACAGGC-3’; Reverse: 5’-ATTTCTGGGCGTGTTGGTGT-3’); *fancd2* (Forward: 5’-CCCTACACTCACCAGGCAAAC-3’; Reverse: 5’-AGCGTTTCAGCTTTCTTGCTATT-3’); *scm2* (Forward: 5’-GCTGAAAGAGAGAAGAAACGCAAA; Reverse: 5’-CTTGCAGAGAGCTCAGACCATC-3’); *rx1* (Forward: 5’-GAGGAACCGGACAACATTCAC-3’; Reverse: 5’-TCATAGCCAGCTCTTZCTCTGC-3’); *gapdh* (Forward: 5’-CTTTGATGCTGATGCTGGA-3’; Reverse: 5’-GAAGAGGGGTTGACAGGTGA-3’).

### WGCNA (Weighted Gene Co-expression Network Analysis)

Network analysis was performed using WGCNA method^17^. Samples used for the workflow were derived from two independent datasets, one from *Nothobranchius furzeri*’s brain, comprehensive of two strains (MZM-04010 and GRZ), six different time points and five replicates per time point ^1^ and the other one from human embryonic stem cells. In particular the second one was obtained from a cerebro-cortical developmental experiment performed on hESC with 9 different time points ^18^.

Network analysis was performed through different steps:

– Setting of the soft threshold, coefficient necessary for the adjacency matrix construction, as shown in the formula:

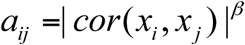
– Adjacency matrix and TOM (Topological Overlap Matrix), defined as:

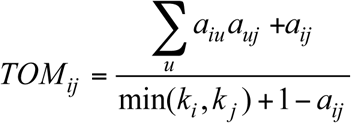
– Hierarchical clustering and modules detection after measuring the module eigengenes; every module is characterized by a color (as the module which has been studied for the analysis, defined by the turquoise color)
Module-trait relationship table construction, as correlation between single gene expression and external trait (in this case aging/development)
– Module membership plot (as correlation between single gene expression and module eigengene): this was done for both the *N. furzeri* and the *H. sapiens* datasets, as described in Figure 5A
– Visualization with Cytoscape software.

Network construction was done in two independent analyses: the first one only on *Nothobranchius furzeri* dataset, the second one using a consensus network obtained matching the two datasets. As soft threshold we chose β=6 for both the analyses to obtain the correspondent adiacency matrix and TOM, and significant modules negatively correlated with *N. furzeri* brain aging were selected. The genes contained in the selected modules were then tested for GO analysis using WebGestalt software, and then visualized using Cytoscape. Finally, the overall module membership of the genes contained in the “turquoise” module (as specified above, and only for the second analysis) was plotted on the ranked genes for both the killifish and the human data. Network analysis was performed using WGCNA method ^17^. Samples used for the workflow were derived from two independent datasets, one from *Nothobranchius furzeri*’s brain, comprehensive of two strains (MZM-04010 and GRZ), six different time points and five replicates per time point^1^ and the other one from human embryonic stem cells. In particular the second one was obtained from a cerebro-cortical developmental experiment performed on hESC with 9 different time points ^18^.

## Author Contribution Statement

V. Naef, S. Monticelli, D. Corsinovi performed all the Xenopus experiments. M.T. Mazzetto and A. Cellerino performed WGCNA (Weighted Gene Co-expression Network Analysis), contributed in the manuscript discussion and writing. M. Ori contributed in conceptualization, provided necessary financial resources, experimental supervision, data analysis and discussion, writing.

## Acknowledgments

We thank Guglielma De Matienzo and Dr. Elena Landi for technical support. This work was supported by funding from University of Pisa (Michela Ori).

